# Long-read sequencing of trios reveals increased germline and postzygotic mutation rates in repetitive DNA

**DOI:** 10.1101/2025.07.18.665621

**Authors:** Michelle D. Noyes, Yang Sui, Youngjun Kwon, Nidhi Koundinya, Isaac Wong, Katherine M. Munson, Kendra Hoekzema, Jennifer Kordosky, Gage H. Garcia, Jordan Knuth, Alexandra P. Lewis, Evan E. Eichler

## Abstract

Long-read sequencing (LRS) has improved sensitivity to discover variation in complex repetitive regions, assign parent-of-origin, and distinguish *de novo* germline from postzygotic mutations (PZMs). Most studies have been limited to population genetic surveys or a few families. We applied three orthogonal sequencing technologies—lIlumina, Oxford Nanopore Technologies, and Pacific Biosciences—to discover and validate *de novo* mutations (DNMs) in 73 children from 42 autism families (157 individuals). Assaying 2.77 Gbp of the human genome using read-based approaches, we discover on average 95 DNMs per transmission (87.5 *de novo* single-nucleotide variants and 7.8 indels), including sex chromosomes. We estimate that LRS increases DNM discovery by 20-40% over previous Illumina-based studies of the same families, and more than doubles the discoverable number of PZMs that emerged early in embryonic development. The strict germline mutation rate is 1.30×10^−8^ substitutions per base pair per generation, strongly driven by the father’s germline (3.95:1), while PZMs increase the rate by 0.23×10^−8^ with a modest but significant bias toward paternal haplotypes (1.15:1). We show that the mutation rate is significantly increased for classes of repetitive DNA, where segmental duplication (SD) mutation shows a dependence on the length and percent identity of the SD. We find that the mutation rate enrichment in repeats occurs predominantly postzygotically as opposed to in the germline, a likely result of faulty DNA repair and interlocus gene conversion.

## INTRODUCTION

*De novo* mutations (DNMs) are variants unique to a child that are absent from its parents. They arise from mutational processes in the parental germline, such as double-stranded break repair or errors in DNA replication^1–5^. In addition, a small number of *de novo* variants arise in the rounds of cell division just after fertilization, early enough in development that they can still be detected in almost all tissues. We term these postzygotic mutations (PZMs) to distinguish them from germline DNMs arising in the parental gametes^6^. The most common type of DNMs are single base-pair substitutions and small (<50 bp) insertions and deletions; previous short-read DNM discovery efforts report an average mutation rate of at least 70 DNMs per individual per generation, over three quarters of which arise in the paternal germline<u>^1,3,4^</u>. PZMs are more challenging to identify and validate, and the PZM rate has been typically estimated to be up to 10% of the germline rate^1,7–10^.

Most studies of *de novo* variation have depended on short-read sequencing (SRS) of parent–child trios or quads to both identify variants and assign them to parental haplotypes. Discovery is typically limited to approximately 84% of the genome where SRS data (150-200 bp) can be reliably mapped^11^. Regions with the largest and most identical blocks of repeats such as segmental duplications (SDs) are typically excluded despite the fact that such regions are predicted to be more mutable^12,13^. Indeed, population genetic and DNM studies on single families have confirmed increased rates of DNM with distinct mutational signatures potentially consistent with the action of interlocus gene conversion but to date there have been few studies from a diversity of families^6,14–16^. In addition to discovery, SRS limits parent-of-origin assignment typically performed using informative single-nucleotide polymorphisms (SNPs), or variants flanking the DNM that can be uniquely traced to one parent or the other. Because the density of SNPs (>1/1200 bp) typically exceeds the average length of SRS (150-200 bp), fewer than 20% of DNMs are routinely phased and assigned to parent-of-origin^3^. Long-read sequencing (LRS) can capture many of these informative SNPs on a single read and can therefore be used to phase more DNMs, which is also useful for distinguishing postzygotic from germline variants due the availability of long-range haplotype information^17,18^.

In this study, we set out to comprehensively identify human germline and postzygotic *de novo* variation across 73 children from 42 families. We leveraged long-read Pacific Biosciences high-fidelity (HiFi) sequencing data derived from blood for variant discovery, and both long-read Oxford Nanopore Technologies (ONT) and short-read Illumina data for validation purposes. These families are part of the Simons Simplex Collection (SSC), and all but one have a proband affected with simplex autism. They have been examined for *de novo* variation before using Illumina whole exome and whole genome sequencing data^12,13,19,20^ but were selected for LRS because no genetic (monogenic or polygenic) cause had been previously identified (Sui et al., companion manuscript). Here, we quantify both germline and postzygotic DNMs in each child. Using LRS, we recover more PZMs than previous Illumina-based studies and assign the parent-of-origin to >97% of all DNMs. The LRS data allow us to evaluate the germline and PZM rates across different genomic regions, quantifying the enrichment of DNMs in hypermutable repetitive sequences, and to compare the mutational signatures.

## RESULTS

### Autosomal *de novo* germline and postzygotic single-nucleotide variants (SNVs)

We examined HiFi data derived from blood and cell lines for a total of 157 samples from 42 families affected with simplex autism (31 quads and 11 trios, n=73 transmissions) for *de novo* variant discovery. While primarily of European ancestry (n=58), the cohort also includes eight children of indigenous American, five of Asian (3 East Asian and 2 South Asian), and two of African ancestry. With T2T-CHM13v2.0 as a reference genome, we applied LRS variant callers to identify SNVs (Methods), selecting those unique to a child for validation with two orthogonal sequencing technologies: ONT and Illumina. For a variant to pass validation, we require that it be observed in a child’s blood-derived HiFi sequence reads in addition to either their ONT or Illumina reads, and that it be absent from both parents across all three types of read data. Additionally, we required that every *de novo* variant is unique to the sample in which it was called; any variant observed in the HiFi reads of unrelated individuals from these families was assumed to be either a segregating, under called parental allele in the parent, or a recurrent sequencing error. Ultimately, 6,030 *de novo* SNVs passed our validation criteria (Methods), for an average of 82.6 SNVs/child.

We classified SNVs as germline or postzygotic in origin initially by haplotype assignment. A variant was considered to be germline if it was present in all the HiFi and ONT reads from its parental haplotype of origin, while a PZM was found only on a fraction of reads from a given haplotype. In other words, PZMs manifest as three or more haplotypes while germline DNMs occur in regions with only two haplotypes where all reads from one parental haplotype share the DNM. Unlike SRS data, the nature of long reads make such distinctions readily possible^6^. In cases of conflict or ambiguity, the small number of remaining variants were classified by examining the allele balance (AB), or the fraction of reads with the *de novo* allele, across all three sequencing platforms (Methods, Figure 1A). We classified 917 SNVs as postzygotic in origin (Figure 1B) while the remaining 5,113 SNVs likely arose in the parental germline (Figure 1C).

**Figure 1:**
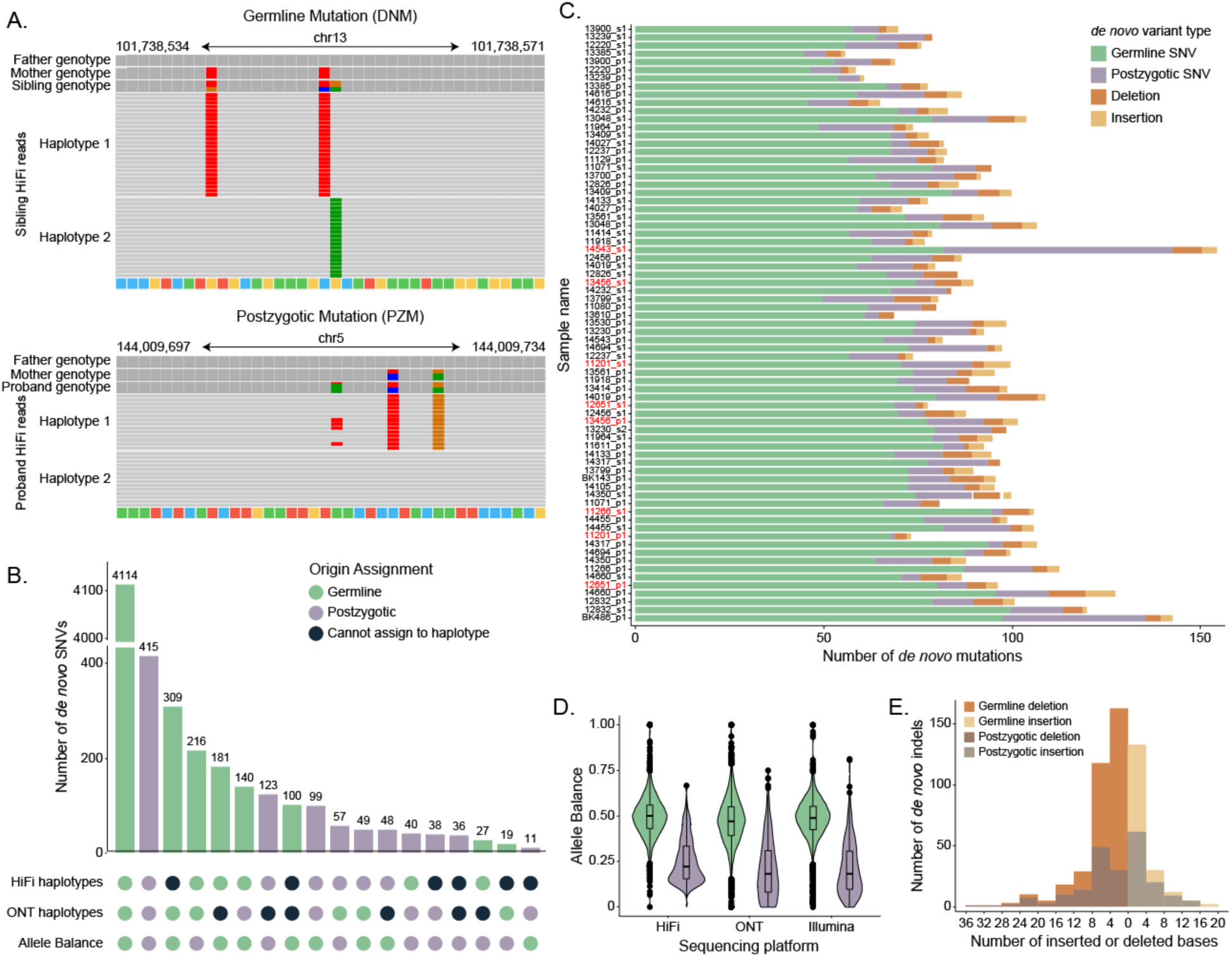
Autosomal SNVs and indels. A. Example IGV views of a germline and postzygotic mutation in HiFi read data. B. The number of autosomal *de novo* germline and postzygotic SNVs, insertions and deletions <50 bp, and tandem repeat mutations observed in each sample. Sibling pairs are grouped by family and highlighted in blue, with the proband above the sibling. C. Upset plot of origin assignment shows concordance between HiFi haplotypes, ONT haplotypes, and allele balance. D. Allele balance distribution for autosomal germline and postzygotic SNVs across PacBio HiFi, Illumina, and ONT read data. E. Distribution of the size of autosomal insertions, deletions, and tandem repeat mutations.

Assessing AB for DNMs across all three sequencing technologies, we calculate the average germline mutation as AB=0.48. In contrast, the average PZM AB is significantly reduced (AB=0.22) (Figure 1D). Although HiFi and Illumina data were derived from blood and ONT data were generated from cell lines to increase scaffolding potential across repeats, we find that AB is remarkably consistent across platforms: 85.7% of PZMs and 89.2% of DNMs do not differ between sequencing platforms (chi-squared test, p-value > 0.05). Notably, a small number (n=11) of germline DNMs have AB=1 across HiFi and at least one other sequencing platform, and all but two of these events fall in repetitive regions (4 in retrotransposable elements, 3 in SDs, and 2 in centromeres).

We were able to assign 98.0% and 96.1% of germline and postzygotic SNVs, respectively, to parental haplotypes. As expected, we observe a significant enrichment of germline mutations on paternal haplotypes (Wilcoxon signed-rank test, p-value = 1.14×10^-13^), with a 3.98:1 paternal:maternal ratio. Surprisingly, PZMs show a modest paternal bias (Wilcoxon signed-rank test, p-value = 0.030), with a 1.15:1 paternal:maternal ratio, although these proportions differ significantly, reflecting their different origins (Z-test (two-sided), p-value < 2×10^-26^). Germline mutations increase in number, by approximately 1.32 and 0.46 additional DNMs per year of paternal and maternal age respectively (linear regression, p-value = 1.24×10^-10^ and 3.32×10^-5^). Parental age has a more modest effect on postzygotic variation, with an additional 0.26 and 0.16 PZMs per year of paternal and maternal age, respectively (p-value = 0.011 and 0.14) (Figure 2C).

**Figure 2:**
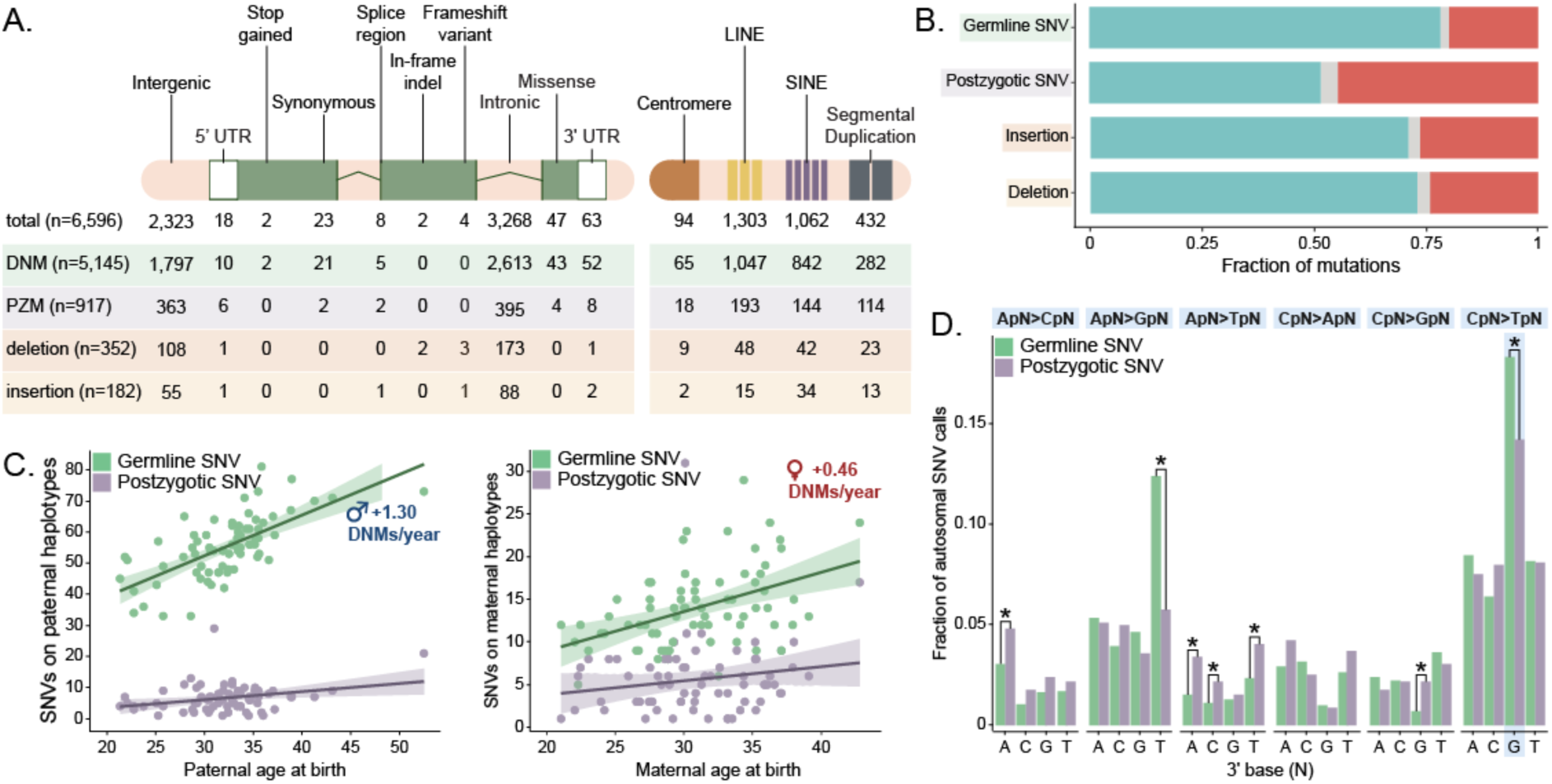
Germline and postzygotic variant comparison. A. Functional annotations of each class of autosomal mutation, as assigned by VEP after lifting variants from T2T-CHM13 to GRCh38. B. Fraction of autosomal mutations assigned to paternal or maternal haplotypes. C. Number of germline SNVs assigned to each parental haplotype as a function of parental age at birth. The slope was defined by linear regression. D. Dinucleotide mutation spectrum of germline and postzygotic SNVs.

Finally, we compared the dinucleotide mutational spectra (Figure 2D) between germline and postzygotic mutations. We find there are 23% fewer postzygotic CpG>TpG mutations relative to germline SNVs. In addition, PZMs are enriched for A>C and A>T substitutions (chi-squared test with Benjamini-Hochberg correction, p-value = 3.06×10^-4^ and 8.05×10^-7^, respectively) and show a marked depletion of A>G substitutions (p-value = 5.36×10^-5^). Notably, we observe an expected transition to transversion ratio (Ti/Tv) of 2.10 for germline DNMs, but PZMs are significantly enriched for transversion mutations, with a Ti/Tv of 1.35 (two-proportions Z-test, p-value = 9.29×10^-10^).

Compared to short-read studies of autosomal SNVs for the same samples, we see a 44% increase compared to Wilfert et al. (n=69 samples in both datasets), a 36% increase compared to An et al. (n=60 samples), and a 30% decrease compared to Turner et al. (n=17 samples)^12,13,20^. Combined, all three studies discovered 1,069 DNMs that were absent from our final callset but could be lifted over to T2T-CHM13v2.0. Approximately 70% of these DNMs were unique to the Turner callset, which we note maximized sensitivity at the expense of specificity. After applying our filtering strategy, we find that 59% and 30% of short-read-only calls appear to be inherited or false positive, respectively, leaving only 166 SNV calls supported by long reads. Of these, only 66 variants passed our full suite of filters, with exactly half excluded because they were observed in two or more unrelated samples. In total, we estimate our callset has omitted 18 false negatives from Wilfert, 8 from An, and 6 from Turner, as well as 34 events from more than one study.

In addition, two samples from this dataset have previously been examined with LRS data, yielding 195 autosomal SNVs that had been validated by ONT and HiFi reads^14^. For the same samples, we identified 196 events, 14.8% (n=29) of which are unique to this study. Of those 29 SNVs, 21 are postzygotic in origin. Conversely, 28 SNVs reported in the previous study are absent from this one. More than a third (n=10) were originally discovered only in GRCh38-aligned reads and were consequently not identified in our T2T-CHM13-aligned data. Of the remaining 18 variants, six were part of mutational clusters that failed our assembly-based validations, five were observed in multiple samples and excluded as errors, four failed validation by T2T-CHM13v2.0-aligned reads, and the remaining three SNVs had conflicting parental haplotype assignments. While the estimated number of DNMs for this family has remained nearly the same, this comparison reveals that we have missed approximately 16 events across these two samples, giving a false negative rate of approximately 7.5%.

### Small *de novo* insertions and deletions

We applied the same criteria used for *de novo* substitutions to identify *de novo* insertion and deletions (indels, <50 bp) with one notable exception; we divided indel calls into two categories: those that were expansions or contractions of short tandem repeats (STRs), and those that were not associated with tandem repeats. Most pure STR mutations were excluded from our indel callset because of the known challenges associated with accurately calling such variants with all three sequencing platforms**^6^**. Outside of STRs, we identified 182 and 351 *de novo* insertions and deletions, respectively, for an average of 7.3 indels/child on the autosomes, with a median indel size of ±4 bp (Figure 1E). This represents a 62% increase over the four indels/child reported by Wilfert et al., a 32% increase over An et al., but a 30% decrease relative to Turner and colleagues’ analysis of the same samples by Illumina sequencing. Overall, we find that 55.7% (n=297) of our indel calls are unique to our HiFi callset and not previously reported in Illumina-based studies. Of the 231 calls observed in Illumina studies that were absent from our final callset, only eight appear to be truly *de novo* across LRS data, while 69.7% (n=161) appeared inherited and the remainder (n=62) were absent from LRS data.

To classify indels as germline or postzygotic in origin, we used a modified version of the same haplotype assignment pipeline. Indels, even those outside of tandem repeats, can be noisier in LRS data, creating multiple possible alleles on a parental haplotype. As a result, indels that are truly germline in origin can appear postzygotic. To account for this noise, we determined an indel was postzygotic if there were more than three reads with an allele that differed from the called *de novo* event. We classified 24.2% of indels (n=129) as PZMs, potentially indicating that significantly more indels are postzygotic in origin compared to the 15% of SNVs that arise postzygotically (chi-squared, p < 2×10^-16^). Germline indels occur on paternal haplotypes at a 3.35:1 ratio, and indels classified as postzygotic occur on paternal haplotypes at a 2:1 ratio (Wilcoxon signed-rank test, p = 8.81×10^-4^). This is a more severe paternal bias than observed for SNVs and is likely due in part to misclassified germline events. There is not a significant relationship between parental age and the size of indel mutations, nor is there a significant relationship between parental age and the number of observed insertions or deletions, arguing strongly that indels within unique regions of the genome arise by distinct mechanisms.

### Sex chromosome DNM

Because of the challenges associated with ploidy differences, we used a modified version of our *de novo* SNV and indel discovery strategies to call variation on the sex chromosomes, treating the females (n=46) and males (n=27) separately. On male X chromosomes, we identify a total of 19 *de novo* SNVs and 5 indels, compared to the 24 SNVs and 8 indels observed on the Y chromosomes (Figure 5A). In female samples, we found a total of 279 SNVs and 24 indels. We determined the origin of female mutations using the same haplotype strategy that we used for the autosomes, and we classified 17.2% (n=48) of female chrX mutations as postzygotic in origin. On the X chromosome, we assigned 166 and 48 germline mutations to paternal and maternal haplotypes, respectively, including 19 SNVs observed in males that must have been inherited from their mothers. We calculate a 3.45:1 paternal:maternal ratio on the X chromosome, completely consistent with the autosomes (two-proportion Z test, p-value = 0.483). We estimate that we were able to discover variation in 96.1% of the female X chromosome, on average (standard deviation [s.d.] = 0.78%) (Figure 5B). As male samples did not inherit an X chromosome from their fathers, we exclude paternal HiFi data when evaluating the male chrX. Conversely, we exclude maternal HiFi data when evaluating chrY. We are able to call on 94.8% of the male X chromosomes (s.d. = 0.80%), which is a small but significant reduction when compared to its female counterpart (t-test, p-value = 9.7×10^−9^). The Y chromosome is highly repetitive, limiting our ability to call in all but 29.3% of the chromosome (s.d. = 0.75%).

### Multinucleotide mutations

Early research of DNMs from parent–child family studies initially reported a small number of clustered mutations sometimes referred to as mutational “storms” or multinucleotide mutations (MnMs)^21,22^. We searched specifically in our callset for such MnMs where two DNMs occurred within 500 bp of each other. We initially identified a total of 106 mutations (n=103 SNVs, n=3 indels), including 23 pairs of *de novo* SNVs immediately adjacent to one another (Figure 3A). However, as part of our initial filtering for DNMs we specifically excluded any mutational clusters where three or more SNVs were located within a sliding window of 1 kbp, effectively removing >1.8 million candidate DNMs from further consideration and biasing against MnMs.

**Figure 3:**
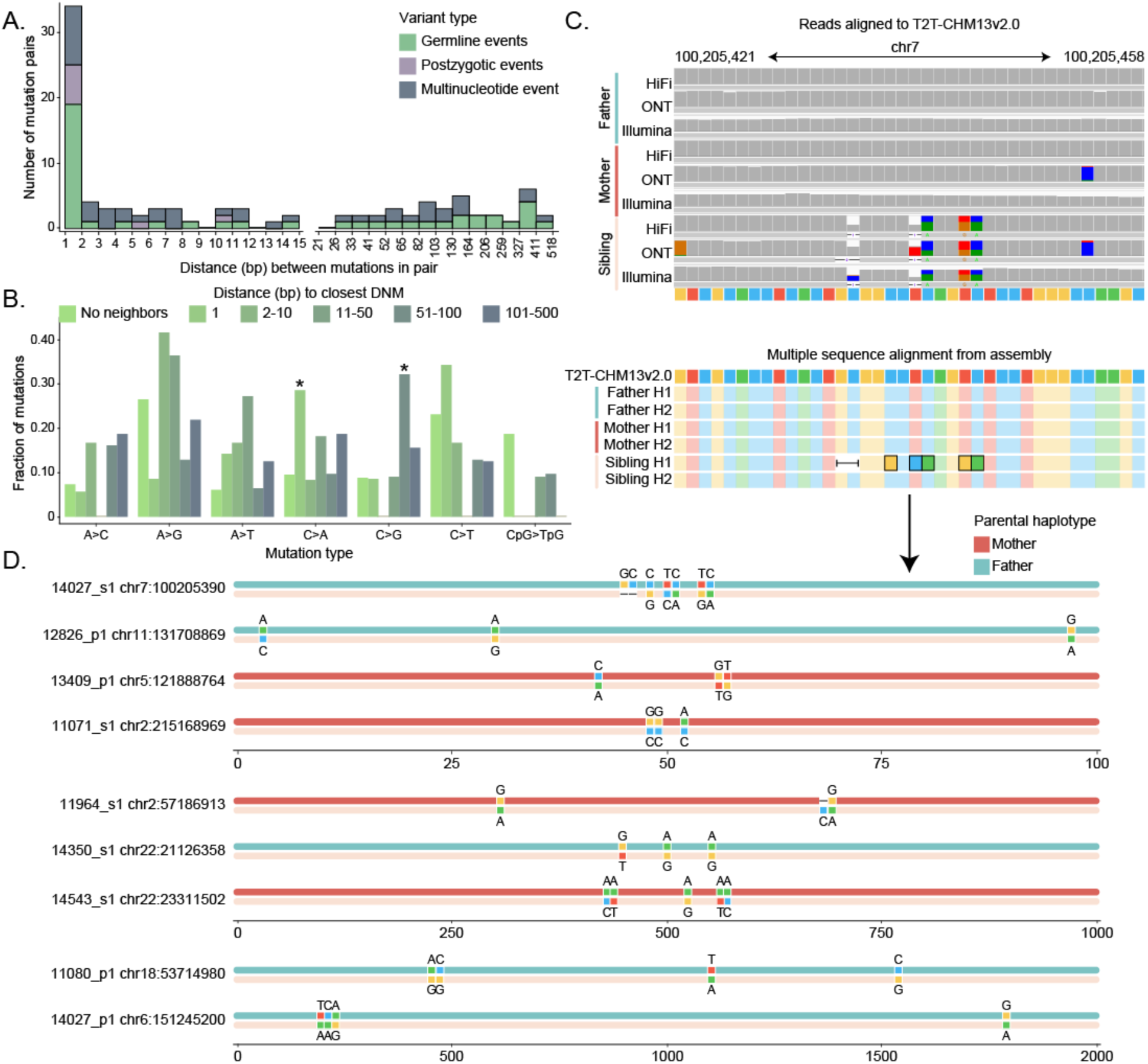
Multinucleotide mutations. A. Distribution of distance between pairs of variants, including SNVs and indels, within 500 bp of each other. B. Mutation spectrum of individual *de novo* mutations (DNMs) and multinucleotide mutations (MnMs), including paired SNV calls within 500 bp of each other. C. An MnM represented in sequencing reads aligned to the T2T-CHM13 reference genome, and in assembled haplotypes from all members of the trio. D. All nine MnMs validated using assembled haplotypes.

We revisited those candidate events applying filters (Methods) essentially excluding >99% of the calls, resulting in a total of 300 candidate single-nucleotide events. For each candidate MnM, we constructed a long-read assembly using hifiasm (**Sui et al., companion**) and compared the parental and child haplotypes. Similar to read-based approaches, we considered a variant to be *bona fide* if it was present in a child’s haplotype and absent from all four assembled parental haplotypes. Finally, we defined variant calls based on the observed mutations in the assembled data—in all cases but one the mutations concurred with the original read-based calls (Figure 3C). This process yielded an additional 44 DNMs, including 8 unclustered SNVs and 1 unclustered indel, and 32 SNVs and 3 indels distributed across 9 clusters (Figure 3D). There was no parent-of-origin bias, as four MnM clusters were observed on maternal haplotypes and five were observed on paternal haplotypes. Seven of these nine clusters corresponded to repeats (4 LINEs, 1 SINE, 1 SD, and 1 low-complexity tandem repeat). Including the paired events from our SNV callset, 50% of the 128 SNVs arising in clusters associate with repetitive DNA, which represents a significant enrichment (Fisher’s exact test, p = 0.0395) compared to unclustered mutations. Further, we find significant differences in the mutational spectra of clustered SNVs when compared to singleton DNMs (Figure 3B). We stratified mutations into six categories based on distance to the nearest SNP and found a depletion of CpG>TpG mutations across all groups of clustered mutations. Although sample sizes are small, we observe a significant enrichment of C>A mutations in adjacent SNPs (chi-squared test with Benjamini-Hochberg correction, p-value = 0.0016) and an enrichment of C>G mutations in SNPs between 51-500 bp apart (p-value = 0.00016).

### DNM rate

In order to determine the DNM rate, we estimate that 91.7% (2.66/2.90 Gbp, s.d. = 23.5 Mbp) (Figure 4a) of the human genome was assayed in this study. Because high LRS mapping quality is used to determine whether a site is callable with our method, we still excluded regions with the highest repeat sequence identity. For example, we can call variants in more than 95% of SDs with sequence identity less than 98%, whereas we can only assess 42% of SDs with over 99% identity. Our read-based approaches perform even worse in centromeres, where we can only assess approximately 6% of higher order repeats and 8% of human satellite sequence. Although we cannot fully examine the variation in these repetitive regions, we are still better equipped to study them than previous Illumina-based *de novo* studies, which typically exclude them completely^2,5,12,13^.

**Figure 4:**
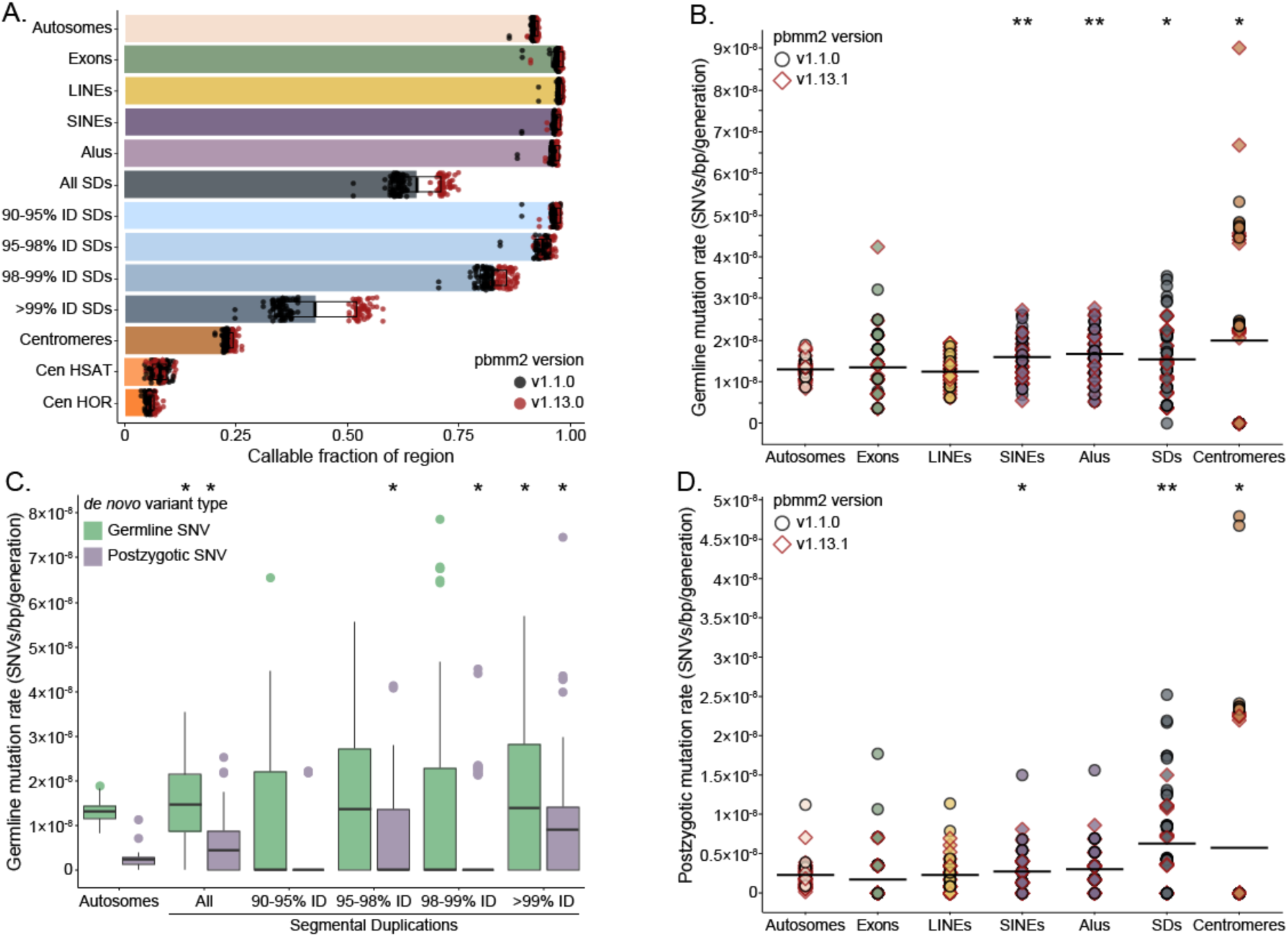
Autosomal SNV mutation rates. A. Mean fraction of callable space in different autosomal regions across all samples in the dataset, with error bars representing 1 standard deviation. B. Distributions of autosomal germline DNM rate for each sample across different genomic regions reveal an enrichment in Alus and segmental duplications (SDs). P-values calculated with t-test and adjusted for multiple testing by Benjamini-Hochberg. C. Autosomal germline *de novo* mutation rates across SDs stratified by percent identity (%ID). D. The same analysis as B, repeated for autosomal postzygotic mutations, finds enrichment in SDs and centromeres.

Thus, limiting to high-confidence regions, we calculate a lower-bound autosomal germline mutation rate of 1.30×10^−8^ substitutions per base pair per generation (95% C.I. 1.27×10^−8^ -1.34×10^−8^) (Figure 4B) and a PZM rate of 2.30×10^−9^ substitutions per base pair per generation (95% C.I. 2.16×10^−9^ -2.46×10^−9^) (Figure 4D). When we restrict our analysis to GENCODEv38 exonic regions of the genome (144 Mbp of sequence, on average 97.2% callable), we find that neither the germline nor postzygotic mutation rate is significantly different from the autosome-wide rate. With respect to the sex chromosomes, we estimate the X chromosome mutation rate to be 1.01×10^−8^ substitutions per base pair per generation in the maternal germline and 2.59×10^−8^ substitutions per base pair per generation in the paternal germline (Figure 5C). This enrichment of mutations in the paternal germline is even more stark on the Y chromosome, where we see a mutation rate of 1.16×10^−7^ substitutions per base pair per generation. Combined with the germline mutations we observed on the autosomes, we calculate the whole-genome mutation rate in the maternal and paternal germline to be 0.52×10^−8^ and 2.09×10^−8^ substitutions per base pair per generation, respectively. The average whole-genome rate is 1.30×10^−8^ substitutions per base pair per generation, which is not significantly different from the autosomal mutation rate.

**Figure 5:**
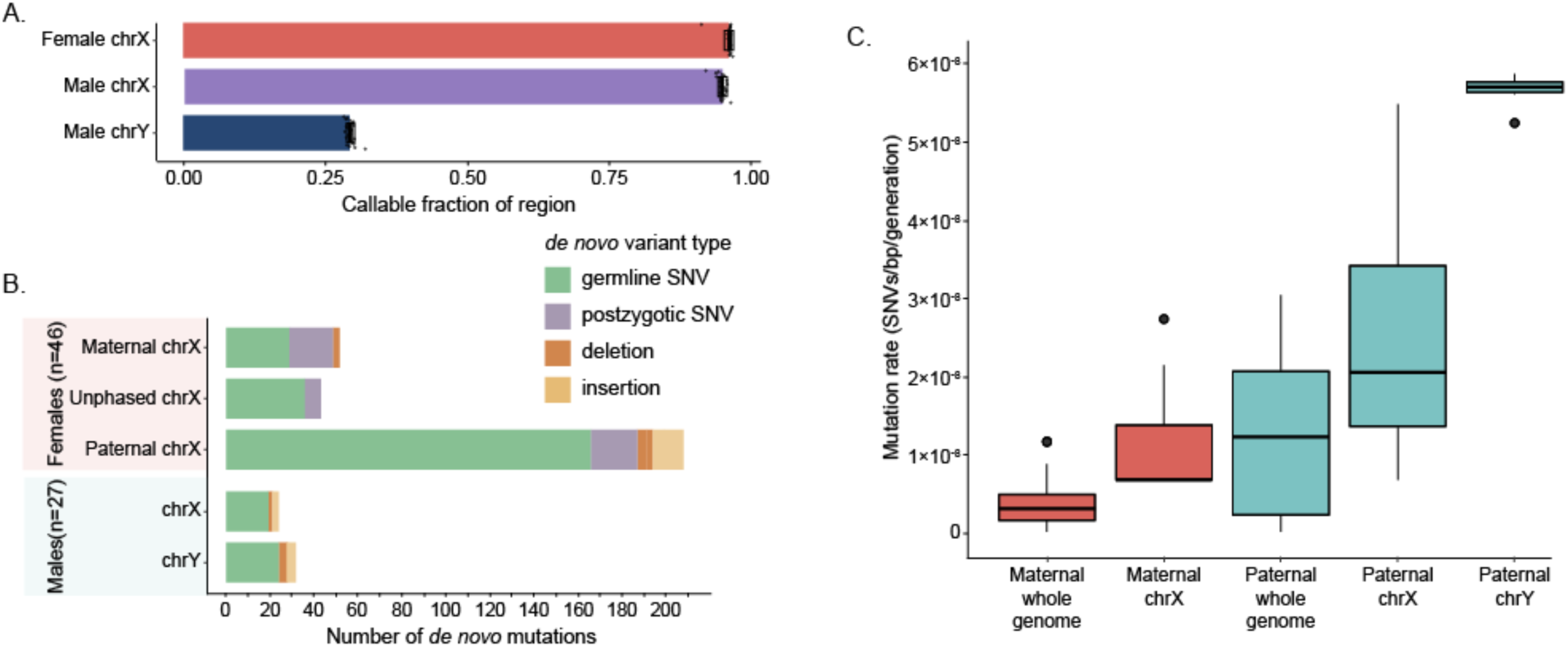
Sex chromosome DNMs. A. Mean fraction of callable sex chromosomes in males and females, with error bars representing 1 standard deviation. B. Number of *de novo* germline and postzygotic SNVs, as well as insertions and deletions, observed on the sex chromosomes. Mutations were phased if possible. C. Distribution of whole-genome and sex chromosome mutation rates calculated using phased germline mutations.

We also contrasted the mutation rate for different classes of repetitive DNA. Notably, we find a germline-specific enrichment in Alus (t-test with Benjamini-Hochberg correction, p-value = 9.93×10^−6^) and SINEs (p-value = 2.35×10^-4^). LINEs, on the other hand, show no enrichment for germline or postzygotic mutations (p-value = 0.34 and 0.96, respectively) relative to the autosomes. In SDs, the germline mutation rate is 1.54×10^−8^ substitutions per base pair per generation (95% C.I. 1.36×10^−8^ -1.72×10^−8^), a significant 18.5% increase over the rate across the autosomes (t-test, p-value = 4.26×10^−2^). This signal is entirely driven by SDs with greater than 99% sequence identity, where the mutation rate is more than double that of the lowest identity duplications (Figure 4C). The PZM rate is more sensitive to sequence identity and is enriched more than twofold in both SDs and centromeres (t-test, p-value = 1.91×10^−6^ and p-value = 3.65×10^−2^). In the case of PZMs, the strongest enrichment is in the highest identity duplications, where we observe an average of 0.85 substitutions per sample (n=62).

## DISCUSSION

Several studies have established that LRS increases the yield of DNM by at least 25% largely by providing better access to repeat-rich regions of the genome^6,14^. In this study of 73 samples from 42 families of diverse ancestry, we identified an average of 95.3 DNM events per child—a 20-40% increase in DNM discovery per sample when compared to earlier short-read studies of the same samples^12,13,20^. Our estimate of the number of SNVs assigned to the germline (83 DNMs/sample) is similar to other reports using LRS (74.5-86) (Tables 1 and 2)^18,23^. Notwithstanding, the number of DNMs per child is significantly less than the 153 DNMs per generation recently reported by Porubsky et al. That study increased sensitivity by applying additional sequencing platforms (Aviti) better equipped to discover indel mutations within homopolymers or STRs, as well as assembly-based approaches to recover DNMs in the most complex regions of the human genome. Thus, while we have increased DNM sensitivity, we still regard our mutation rates as a lower bound, significantly underestimating the indel mutation rate, especially in STRs and overall in more complex regions of the genome.

Another major advantage of LRS is the ability to phase variants assigning them unambiguously to parental haplotypes. Without pedigree information, short-read data can typically be used to phase up to 20% of DNM calls^2,3^, but we were able to phase 97.7% and 97.0% of autosomal SNVs and indels, respectively. In addition, the phasing data can be used to classify DNMs arising in the germline versus the early embryo. In our study we estimate that 15.1% of SNVs (12.6 DNMs per child) are postzygotic in origin, almost doubling earlier estimates of 4-10%^1,7,9,10^. Although it is possible that these PZMs represent somatic mutations that arose and clonally expanded in the blood, this higher estimate is consistent with our recent multigenerational LRS sequence assembly analysis where we showed that ∼60% of the PZMs are transmitted to the next generation^6^. That analysis confirms that a significant fraction of such PZMs are present in the germline (in addition to the blood) and suggests at least two-thirds of the PZMs discovered here are likely early embryonic mutations.

PZMs show unique mutational signatures and biases compared to germline mutations, likely due to the different environment in which they arose. PZMs are depleted for CpG-associated mutations and are enriched for transversions when compared to those classified as germline (postzygotic Ti/Tv of 1.3 vs. germline Ti/Tv of 2.1). We find that 80% (3.95:1) of germline mutations occur in the paternal germline, where an additional 1.3 SNVs arise with every passing year, in contrast to PZMs, which are almost equally distributed among parental haplotypes (53% of PZMs arise on paternal haplotypes). This modest but significant paternal bias for PZMs (1.15:1) with evidence of a slight paternal age effect of 0.26 PZMs per year could suggest germline involvement. One possibility may be single-stranded DNA damage exists as heteroduplex in the sperm that cannot be adequately repaired until after conception. Even after fertilization, DNA transcription does not begin until the 4-8 cell stage^24^, so some lesions may persist, constrained to the repair machinery available in the oocyte^25^. Coupled with error-prone early cell divisions^26,27^, the embryo may preferentially turn to repair mechanisms like allelic and interlocus gene conversion (IGC) to correct errors.

Based on the 91% of the callable genome, we calculate a germline mutation rate of 1.30×10^−8^ substitutions per base pair per generation. Excluding PZMs, we predict the mutation rate for 30-year-old parents to be 1.25×10^−8^, which is very close, on first blush, to estimates of 1.21-1.30×10^−8^ made by SRS- and LRS-based studies^1,2,4,6,23,28^ (Supplemental Table 1). We estimate, however, that PZMs contribute an additional 0.23×10^−8^ substitutions per base pair per generation for a combined germline and PZM DNM rate of 1.53×10^−8^—a marked increase over short-read-based estimates. Adjusting for PZMs, the germline mutation rates from previous studies would more likely fall into the range of 1.1-1.21×10^−8^.

Consistent with our study of a multigenerational family^6^, the mutation rate is elevated in repetitive DNA but not uniformly so. We find significant increases in DNM for SINEs and SDs but not LINE repeats, perhaps due to the decreased CG content for the latter and the suspected role of IGC in elevating the mutation rate of these regions^29–31^. Both germline mutations and PZMs are significantly enriched in SDs and the acceleration increases the longer and more identical the repeats become (Figure 4C). Specifically, the germline mutation rate for SDs >99% sequence identity is 30% higher while the postzygotic rate is over fourfold higher. These findings suggest that SDs are particularly prone to substitution mutations early in embryogenesis. Although the strict germline mutation rate increase (18.5%) is more modest than we might expect based on previous population genetic estimates (which predicted an increase of 60%)^31^, if we combine both the germline and postzygotic rate, we estimate an overall 42% increase in mutation over SDs. The fact that DNMs in SDs are significantly depleted in CpG substitutions and show an excess of transversions once again points to a role for IGC in driving some of this acceleration^31^. What’s more, these signatures are also consistent with the types of PZMs observed in SDs.

Another potential result of IGC and repeat-associated mutation is clustered *de novo* MnMs. We identified nine such MnM clusters that were supported by both read-based and assembly alignments. Six of the mutation clusters mapped to repeats, including LINEs (n=4), SINEs (n=1), and SDs (n=1). We hypothesize that some of these MnMs are likely the result of faulty repair with polymerase ζ, which preferentially creates paired single-nucleotide substitutions, and in fact 9 of our 23 tandem SNV mutations are GC>AA or GG>TT substitutions, a hallmark of this polymerase^22,32^. Notably, four of our MnM clusters have an exact sequence match to the *de novo* allele in a paralogous repeat mapping elsewhere in a parental genome, suggesting these sequences served as the original template for the IGC event^33^. Comparatively, only 6% of germline SNVs and 8% of postzygotic SNVs are gene conversion candidates. The plurality of conversion candidates are in LINEs (44%), with an additional 20% each in Alus and SDs. While it is likely that many conversion candidates did not actually arise from IGC events, IGC clearly contributes to the mutational landscape of these repetitive regions.

Despite the increased mutation rates we report, additional DNMs remain to be discovered. While we were able to assess small variants in approximately 91% of the genome, the remaining 9% is among the most mutable^6^. Owing to the unreliability of Illumina and LRS-based variant calls, for example, we excluded some of the most complex tandem repeat insertions and deletions within STRs and VNTRs from this analysis. We only analyzed approximately 6% of centromeric alpha-satellite DNA and excluded over three quarters of the Y chromosome where mutation rates are known to be more than order of magnitude higher^6,34^. We also did not assess large-scale structural variation because most *de novo* SVs map to complex regions ill-suited for characterization by read-based alignments^6^. Accessing these regions and these particular classes of mutation will require assembly-based approaches which, in turn, require sequencing of higher molecular weight DNA, not as readily derived from retrospective clinical research material. A dedicated effort to sequence genomes telomere-to-telomere from many more diverse families should be undertaken to better understand patterns, rates, and biases of DNM among humans.

## Acknowledgements

This work was supported, in part, by the US National Institutes of Health (NIH R01MH101221 to E.E.E.) and the Simons Foundation (SFARI #810018EE to E.E.E.). E.E.E. is an investigator of the Howard Hughes Medical Institute. We thank Tonia Brown for assistance in editing this manuscript.

This article is subject to HHMI’s Open Access to Publications policy. HHMI lab heads have previously granted a nonexclusive CC BY 4.0 license to the public and a sublicensable license to HHMI in their research articles. Pursuant to those licenses, the author-accepted manuscript of this article can be made freely available under a CC BY 4.0 license immediately upon publication.

## METHODS

### Illumina sequencing

We used previously published Illumina data generated by the NYGC for the SSC^12–14^. Briefly, each sample was sequenced on the Illumina X Ten platform using 1 μg of blood-derived DNA with an Illumina PCR-free library protocol.

### HiFi sequencing

We used the same method as described in Sui et al. (companion piece) to generate HiFi data from blood- and lymphoblastoid cell line-derived DNA.

### ONT sequencing

ONT data were generated from DNA extracted from lymphoblastoid cell lines using a modified Gentra Puregene protocol. Libraries were constructed using the Ligation Sequencing Kit (ONT, SQK-LSK110) with modifications to the manufacturer’s protocol. The libraries were loaded onto a primed FLO-PRO002 R9.4.1 flow cell for sequencing on the PromethION, with two nuclease washes and reloads after 24 and 48 hours of sequencing.

### Alignment to the reference genome

We selected T2T-CHM13v2.0 as our reference genome, as it allows us to align reads to repetitive regions that were not represented in the previous reference. For all females, we masked the Y chromosome from the reference before alignment, and for all males we masked the pseudoautosomal regions of the Y, following the guidance from Rhie et al.^35^ Illumina data were aligned using BWA-MEM v0.7.17.^36^ We aligned both HiFi and ONT data using minimap2^37^, with the help of the pbmm2 v1.13.1 (https://github.com/PacificBiosciences/pbmm2) wrapper for handling the HiFi data. It is important to note that we aligned the first batch of processed samples (n=42) using pbmm2v1.1.0 and the second batch of samples (n=73) using pbmm2v.1.13.0. The newer version of pbmm2 performed notably better in higher sequence identity regions, but there was no difference in the average number of DNM calls found across both batches of samples.

### *de novo* SNV discovery and validation on the autosomes

Variant calling was performed using aligned HiFi data and two variant callers, GATK HaplotypeCaller v4.3.0.0^38^ and DeepVariant v1.4.0^39^, following the same filtering strategy outlined in Noyes et al. 2022^14^. For each caller, we naively identified candidate *de novo* events by selecting any variant where both parents were homozygous for the reference allele and the child had at least one alternate allele. We took the union of both candidate *de novo* callsets, retaining only variant calls where the child’s genotype quality was at least 20, resulting in an initial callset of 2,159,552 SNVs across all 73 samples. To eliminate runs of candidate *de novo* events that were actually the result of a dropped haplotype in one parent, we eliminated any regions where three or more SNVs were found in a sliding 1 kbp window, removing a total of 1,837,202 candidate events from our callset.

Next, we examined the HiFi, ONT, and Illumina reads that spanned each candidate variant in a child and both parents. For a child’s HiFi read to be considered, it had to be derived from blood data, but both parental blood and cell line reads were retained (all ONT data were derived from cell lines; all Illumina data from blood). Long reads with mapping quality <59 were excluded (we did not filter short reads on the basis of mapping quality). We partitioned reads into three categories based on the base quality (probability that a base was correctly called) at the site of the variant: reads with base quality >20 (high quality), reads with base quality between 10 and 20 (low quality), and reads with quality <10, which were discarded. For each sequencing platform, we counted the number of reads that supported the reference and alternate alleles in both parents and the child and used them to determine whether a variant was truly *de novo* or inherited. For HiFi and Illumina data, we required that each parent have fewer than one high-quality or two low-quality reads with the *de novo* allele, and the child have at least one read with the *de novo* allele. Since ONT is slightly less accurate, we required fewer than two high-quality or three low-quality reads with the *de novo* allele. Once each variant was examined in each platform, we combined the validations, determining that a variant was inherited if it looked inherited in at least one platform, and truly *de novo* if it was supported in at least two platforms. Across all samples, 14,187 candidate mutations passed this initial round of filtering.

We returned to the aligned HiFi reads for every sample in the dataset (parents and children), checking every candidate *de novo* allele to see if it was represented in more than one sample, removing approximately 7,000 variants that we determined to be recurrent errors. If a variant was not present in a tandem repeat (TR), we required that it be unique to the child it was identified in, and if the variant was in a TR, we allowed it to be observed in one unrelated sample. In addition, variants in TRs had to have an average allele balance (AB) greater than 0.05 across all platforms. We then removed low-quality variants, that had dubious support across two or more platforms (typically noisy parental data with alternate alleles different from the *de novo* allele). We excluded variants in regions flagged by RepeatMasker that failed AB filters (0.1 if it was also in a TR, 0.08 if not)^40^. Lastly, we removed variants in or adjacent to homopolymers that involved the homopolymer subunit (i.e., an A to T substitution on the edge of an A homopolymer), as those variants are typically sequencing artifacts and difficult to validate across all sequencing platforms. We validated a total of 6,070 variants that we then assigned to haplotypes. A final 40 variants could not be uniquely assigned to one parent and were excluded, resulting in a final callset of 6,030 autosomal SNVs.

### *de novo* indel discovery and validation on the autosomes

We generated candidate indel callsets using the same combined GATK and DeepVariant callset, naively selecting for any insertions or deletions present in a child and absent from its parents. We divided these calls into two categories: TR mutations—where one or more subunits were added or subtracted, and indels—that either did not involve a perfect TR motif or did not overlap with a TR (n=616,873). We applied a similar read-based validation strategy for indels as we did for SNVs. We required that HiFi, ONT, and Illumina reads have mapping quality of 60 (the highest possible) and that they fully span the variant site, with at least 10 bp of flanking sequence before and after, to ensure that we captured the full allele.

We excluded variants in TRs. To validate indels outside of TRs, we examined child and parental data across all three sequencing platforms, counting the number of *de novo* alleles. If a child had a sibling in our dataset, we also examined the sibling’s read data. We considered a variant to be inherited if one parent or the sibling had a read supporting the *de novo* allele in any platform. We deemed a variant to be truly *de novo* if we observed the *de novo* allele in the child in both HiFi and Illumina data, resulting in a total callset of 596 *de novo* indels. Finally, we visually inspected every indel call in IGV, removing any indels with several alternate alleles. Many of these sites were located in homopolymers or repetitive sequence, resulting in a diversity of alleles and often obscuring the inheritance pattern, resulting in false positive calls. After manual inspection, our final callset was composed of 533 indels.

### Multinucleotide mutation (MnM) discovery

To identify MnMs, we began with the excluded 1,837,202 calls from the SNV validation pipeline. We removed the clustered mutation filter and applied the same validation steps as for unclustered SNVs. Only 15,713 mutations remained after the initial three-platform validation, and 300 passed every filter. We phased these 300 SNVs, excluding 10% (n=30) because they were found on both parental haplotypes. For the remaining 270 SNVs, we examined parent and child assemblies generated with hifiasm (Sui et al., companion)^41^.

We defined MnM regions by examining GATK VCFs, selecting any SNVs within a 5 kbp sliding window around the validated event. We then subset these regions from parent and child assemblies, only retaining variants when all four parental haplotypes and both child haplotypes were fully assembled in the 50 kbp surrounding the clustered event. We used MAFFT^42^ to generate a multiple sequence alignment of all six assembled sequences, as well as the corresponding sequence from T2T-CHM13v2.0. We retained any variants that were unique to the a child’s haplotype and excluded cases in which the surrounding sequence had excessive mismatches surrounding the region, a phenomenon we commonly observed in misaligned or misassembled regions such as centromeres. This process yielded 44 DNMs, 9 unclustered events (8 SNVs and 1 indel), and 9 clustered events. In cases where the assembly-based variants disagreed with the GATK-called variants, we deferred to the assembly-based results (Figure 3C).

### Sex chromosome *de novo* discovery

To identify variation, we used ploidy-aware GATK HaplotypeCaller v4.3.0.0, treating the female chromosome X as diploid, and the male sex chromosomes as haploid. Females and males from each family were jointly genotyped separately, and variant calls were filtered using the same parameters as autosomal variants.

For female children, we naively identified *de novo* variation by selecting sites that were homozygous reference in the mother and hemizygous reference in the father, and the child had an alternate allele, identifying 29,195 SNVs and 17,772 indels. We excluded 21,838 SNVs that were in clusters of three or more within 1 kbp, then used most of the SNV filtering strategy that we applied to autosomal variants, examining HiFi, ONT, and Illumina reads to ensure each variant was unique to the child in which it was identified. We used the same filtering parameters for sites in TRs but did not filter based on RepeatMasker or homopolymer annotations^40^, as few sites were in such regions. In total, 283 SNVs on female X chromosomes passed our validations. After assigning these variants to parental haplotypes, four had conflicting parental information and were excluded from our final callset of 279 SNVs. We used the same autosomal indel validation pipeline for non-TR variants, resulting in a final callset of 26 indels on female X chromosomes.

For male children, we treated X and Y chromosome variation separately, excluding the pseudoautosomal regions on both. We naively identified variants on the X chromosome by comparing a child to his mother, selecting any sites where the child had a different allele (not required to be reference or alternate). Conversely, we selected sites on the Y chromosome where a child had a different allele than his father. We identified 21,107 SNVs and 78,939 indels on male X chromosomes, and 100,225 SNVs and 8,215 indels on male Y chromosomes. To validate SNV calls on the male X chromosome, we applied the same filtering strategy as the female X, first evaluating HiFi, ONT, and Illumina sequencing data to ensure that a variant was unique to a child, and then filtering sites in TRs, resulting in a total of 19 SNVs on male X chromosomes. For SNV calls on the Y chromosome, we simply checked a child and the father’s sequencing data across all three platforms to ensure that the variant was not present in the father, resulting in 140 SNVs. We manually validated all 140 SNVs in IGV and excluded any that had low sequencing depth (fewer than 3 reads) and extensive local variation, resulting in a final callset of 24 SNVs on male Y chromosomes. We used the same indel filtering strategy for male sex chromosome variants as for female, except we only examined maternal sequencing data for X and paternal data for Y chromosome variants. In total, we found three indels on male X and eight indels on male Y chromosomes.

### Phasing

For every DNM, we identified informative SNPs within an 80 kbp window centered at the mutation site based on variant calls from our GATK4 run using HiFi data. An informative SNP is defined as any SNP whose origin can unambiguously be assigned to one parent: for example, a site where one parent is 0/0, the other parent is 0/1 or 1/1 and their child is 0/1. We then examined the HiFi read data for the child, examining every read derived from blood DNA that passed our read filters (mapping quality ≥ 59 and base quality ≥ 20 at the site of the DNM). We assigned each read to a maternal or paternal haplotype by calculating an inheritance score based on the presence of tagging SNPs. We gave tagging SNPs a value of ±1 depending on whether they were inherited from the mother or father, and then took the average of these values, inversely weighted by each SNP’s distance from the DNM site. A negative inheritance score indicated a paternally inherited read, and positive score indicated a maternally inherited read. If all the reads with the *de novo* allele could be assigned to one parent, we assigned them as the parent-of-origin. If the *de novo* allele was present on both paternal and maternal haplotypes, it was left unphased. Using this method, we were able to phase 90.7% of all SNVs (n=5,509), while the remaining 561 had no tagging SNPs or ambiguous parental data.

DNM origin was also evaluated using ONT reads. We applied the same read-filtering steps, with the caveat that all ONT data were derived from cell lines, and calculated inheritance scores based on informative SNPs. For each SNP, we compared the HiFi and ONT haplotype assignments, preferentially selecting the HiFi assignment in cases of disagreement (n=8) between the two sequencing platforms. With the ONT data, we were able to phase an additional 417 DNMs, leaving just 3% of our DNMs (n=136) unphased. In cases where tagging SNPs were present, but neither ONT nor HiFi data could unambiguously phase a *de novo* variant (n=48), we assumed the variant was due to sequencing error and excluded it from the final DNM callset.

### Assessment of postzygotic mutation

Using the filtered and parentally assigned HiFi and ONT reads from our phasing pipeline, we counted the number of reference and alternate alleles derived from each parent and determined that a DNM was postzygotic in origin if we detected at least two reads with the reference and one read with the alternate allele assigned to one parent’s haplotype.

We also predicted whether a mutation was likely to be germline or postzygotic based on the new variant’s AB across HiFi, ONT, and Illumina sequencing data. First, we filtered reads according to the same parameters used in the phasing pipeline, restricting HiFi data to only blood-derived reads, and counted the number of reads with the reference and alternate alleles. We used these counts to calculate AB for each platform, and then used a chi-squared test (using chi2_contingency from the python package scipy.stats) to determine whether the three AB values were concordant. In cases where the AB was not concordant, we could not confidently predict whether the variant was postzygotic, so it was supposed to be germline. If the AB was concordant across platforms, we pooled the reference and alternate allele counts and tested whether the total AB was significantly less than 0.5 using a binomial test (binomtest from scipy.stats). Any variants with significantly low AB were predicted to be postzygotic in origin.

To make the final determination of mutation origin, we combined results from the HiFi and ONT haplotypes and AB-based predictions. In cases where HiFi and ONT haplotypes disagreed (n=286), we used the origin assignment that matched our AB prediction, and in cases where HiFi and ONT haplotypes were ambiguous (n=184), we used the AB prediction. In total, we determined that 917 *de novo* SNVs were likely postzygotic in origin, and 5,145 arose in the parental germline.

### Callable genome and mutation rate calculation

To determine where we were able to identify *de novo* variation in the genome, we assessed HiFi data for every trio. We first used GATK HaplotypeCaller v4.3.0.0 with the option “ERC BP_RESOLUTION” in order to generate a genotype call at every site in the genome. Only sites where both parents were genotyped as homozygous reference (0/0) were considered callable, as sites with a parental alternate allele were excluded from our *de novo* discovery pipeline. We then examined the HiFi reads from a sample and its parents, restricting to only primary alignments with mapping quality of at least 59. For children, we only considered HiFi reads derived from blood, but we considered blood and cell line data for parents. We counted the number of reads with a base of at least a quality score of 20 at every site in the genome, and then combined this information with our variant calls. A site was deemed callable if both parents and the child each had at least one high-quality read with a high-quality base call. We observed an average of 2.66Gbp (out of 2.90 Gbp, s.d. = 24.9 Mbp) such sites across the autosomes. For female children, the callable chromosome X was determined the same way, whereas for the male children, we only considered the mom’s HiFi data when examining the X chromosome and the dad’s HiFi data when examining the Y chromosome. In addition, male sex chromosomes were not restricted to sites where both parents were genotyped as reference—each parent was allowed to carry an alternate allele.

It is important to note that we processed our HiFi data in two separate batches, aligning them with two versions of pbmm2 (v1.1.0 and v.1.13.1). Due to improvements in alignment quality over repetitive regions, samples aligned with the newer version have an average of 24 Mbp of additional callable space across the autosomes, with the most notable gains in high-identity SDs. We did not observe an increase in DNM calls in samples aligned with the newer version of pbmm2.

We calculated the germline autosomal mutation rate for every sample by dividing the number of germline autosomal DNMs by twice the number of base pairs we determined to be callable. For PZMs, we used the same denominator. In females, the amount of callable sex chromosomes was defined as twice the number of callable bases on the X chromosome, and in males it was defined as the sum of the callable bases on the X and Y chromosomes. For each feature-specific mutation rate (such as SDs), we intersected both a sample’s *de novo* SNVs and the sample’s callable regions with coordinates of the relevant feature. We then calculated the mutation rate by dividing the number of SNVs in the region by the amount of callable space in the region.

