## Supplemental Note (Supplementary Figures 1-5) for "Long-read sequencing of trios reveals increased germline and postzygotic mutation rates in repetitive DNA"

SUPPLEMENT

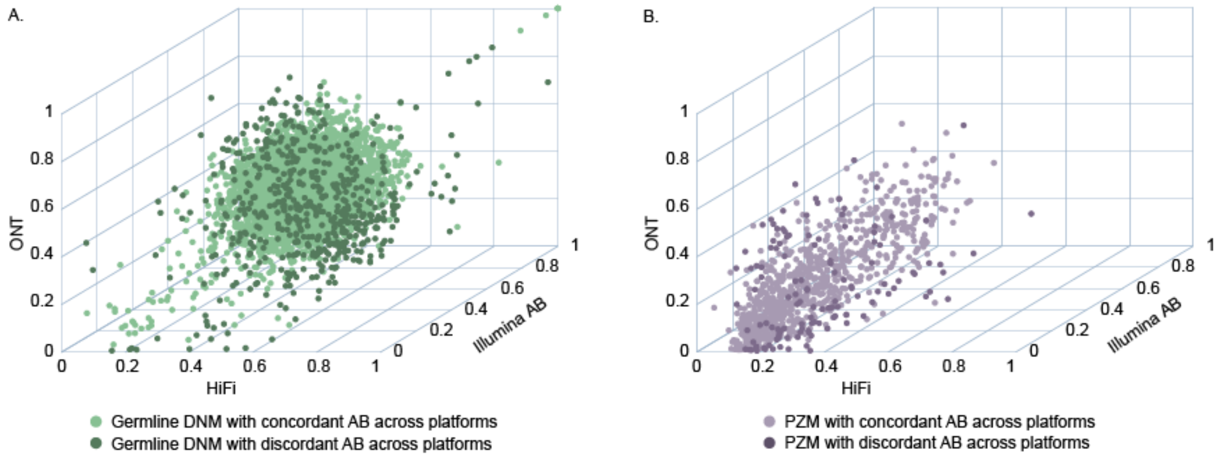


Figure S1: Allele balance scatterplots

A. Allele balance (AB) of germline DNMs across all three platforms for variants with concordant and discordant AB by chi-squared shows that most variants have AB around 0.5. B. PZMs tend to have low AB across platforms.

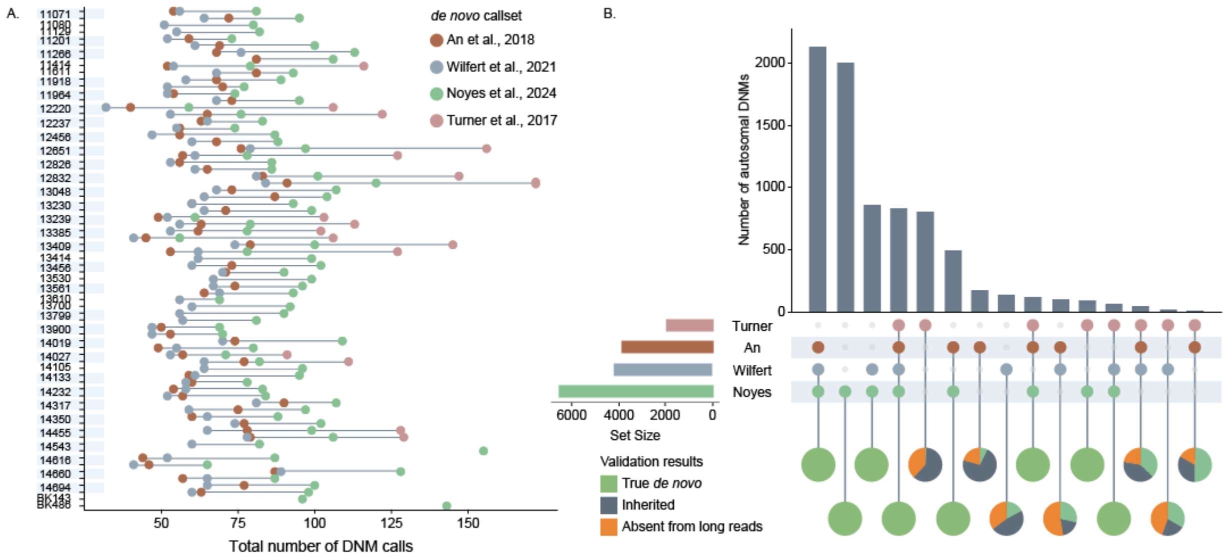


Figure S2: Comparison to other studies

A. We identified more DNMs per sample than most previous Illumina-based studies, with the exception of Turner et al. B. DNMs identified by multiple studies have the highest true positive rates, while DNMs exclusive to a single study tend to be false positives.

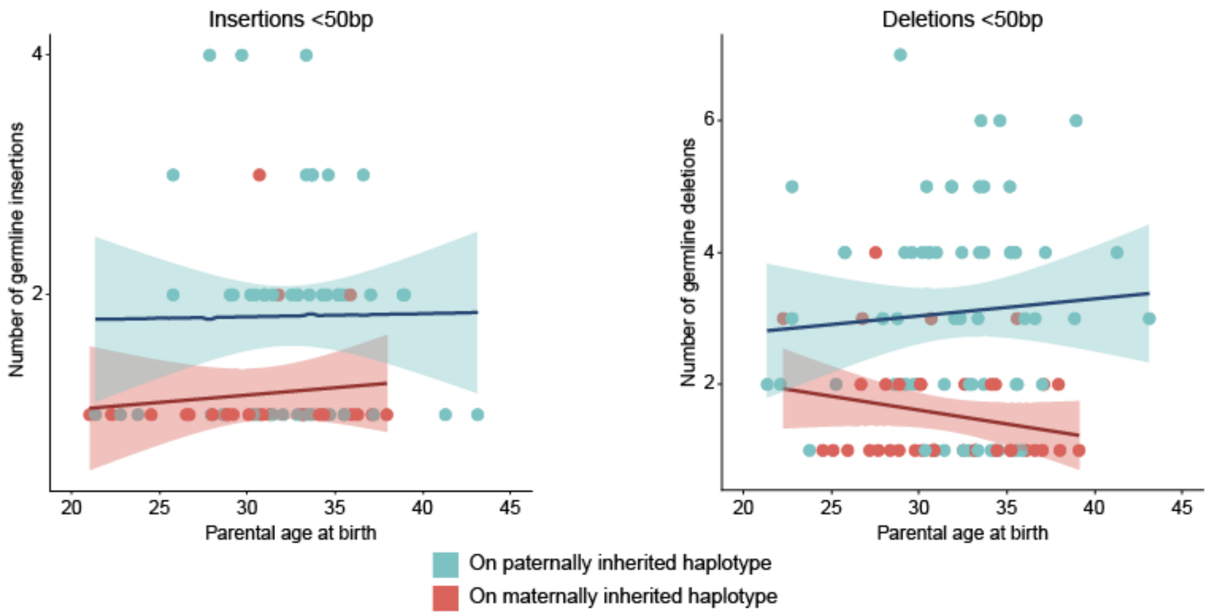


Figure S3: Indel parental age effect
Neither insertions nor deletions are significantly correlated with year of parental age.

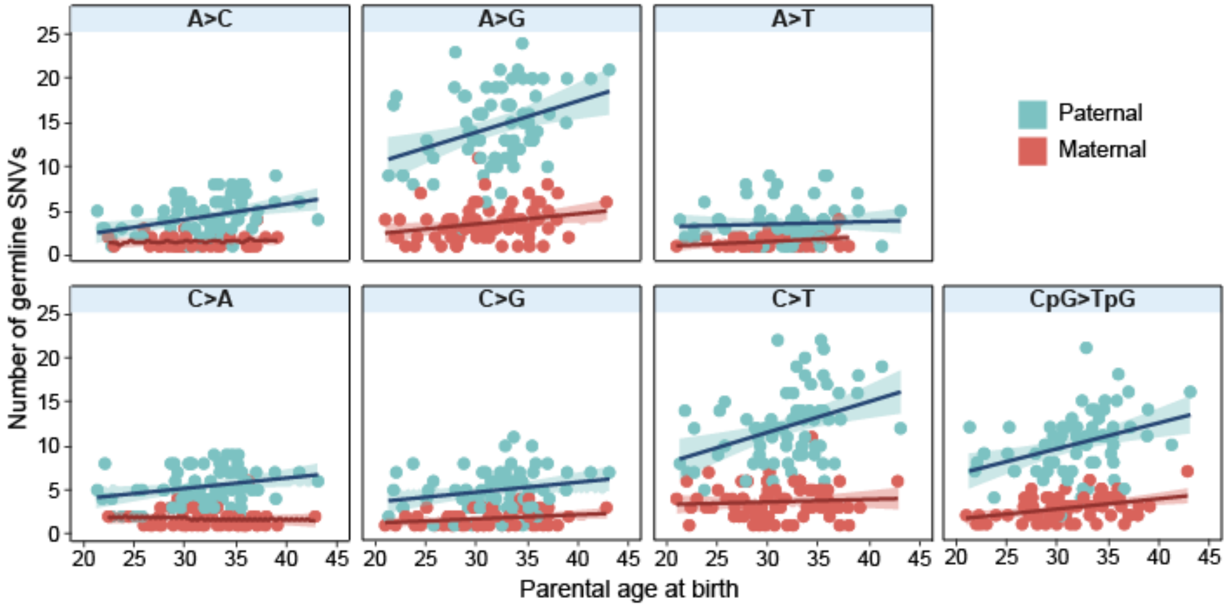


Figure S4: Parental age effect by mutation class
The number of paternal A>C, A>G, C>A, and CpG>TpG mutations are all significantly correlated with paternal age. For maternal mutations, there is a significant maternal age effect on A>G and CpG>TpG mutations.

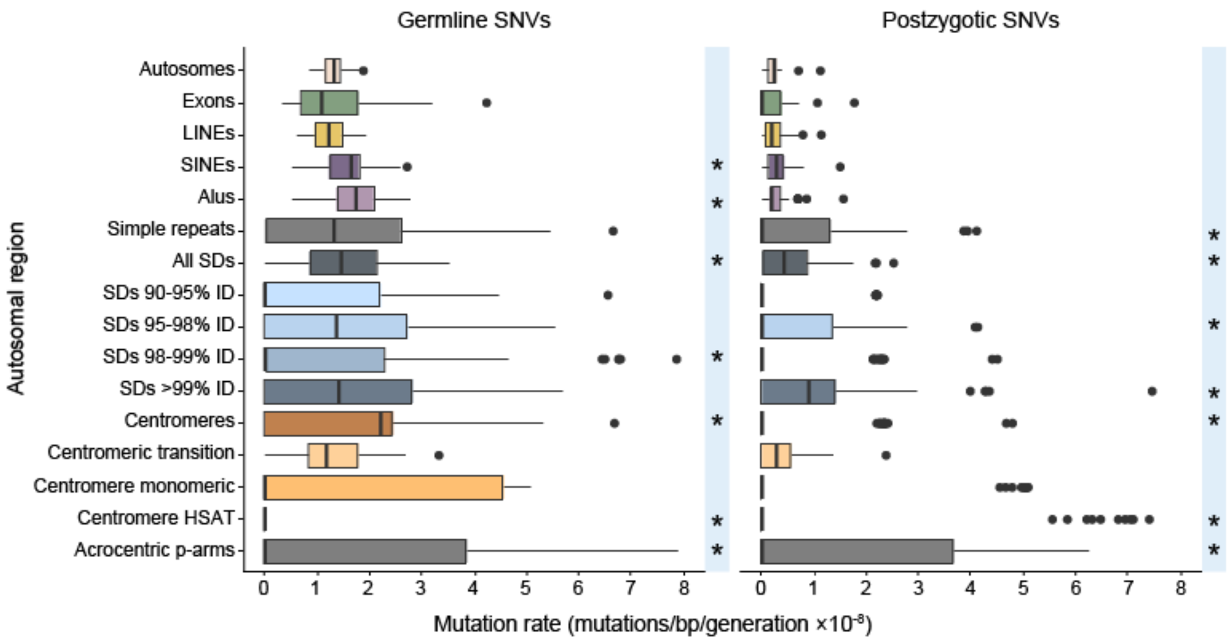


Figure S5: Mutation rates in segmental duplications and centromeric regions

The germline and postzygotic mutation rates are significantly enriched in the highest-identity segmental duplications and acrocentric p-arms.
